# The Virtual Child Brain: Modeling Neuromaturational Trajectories

**DOI:** 10.64898/2026.07.07.737052

**Authors:** Karin Westin, Leon Martin, Marius Pille, Michael Schirner, Petra Ritter

## Abstract

**Introduction:** Understanding the mechanisms of human neuromaturation constitutes one of the fundamental questions of neuroscience. While it is well described that large-scale brain maturation is initiated within sensorimotor brain regions and progresses to associative cortex, the underlying developmental neurobiology remains to be fully characterized. Animal models have indicated that cortical inhibitory upregulation might be a driver of neurodevelopment. To investigate the hypothesis that cortical inhibitory upregulation plays a similar role in human neuromaturation, we developed a The Virtual Brain (TVB) based computational model (TVB-Child) to explore potential mechanisms of human neurodevelopment.

**Material and method:** We created neurodevelopmental dynamic brain network models capturing neurobiological maturation by using the large-scale brain simulator TVB and fitting brain network models to developmental functional MRI (fMRI) from the Human Connectome Project-Development (HCP-D) data set with 640 subjects with an age range of 6-21 years. Age-dependent trajectories in the fMRI data set were first analyzed by combined group-ICA/Dual Regression extracting subject-specific resting-state networks (RSN). Maturational topographical and topological redistribution of these networks were analyzed by linear and non-linear regression of RSN size and degree and strength centrality. Brain network models were fitted to the fMRI functional connectivity obtained from the HCP-D data set. Hypothesizing that cortical inhibition is a driver of neuromaturation, we analyzed spatiotemporal inhibition parameter gradients in the dynamic brain network model for the hypothesized significant correlations with fMRI RSN maturational trajectories.

**Results:** While during development frontoparietal (FP) and default mode network (DMN) grew and exhibited an increase in both degree and strength centrality, becoming dominant network hubs, the attention network underwent network pruning with a decrease in size and node degree. The primary sensory network changed little. For the fitted brain network models, we obtained a high degree of reproduction with correlation coefficients between empirical and simulated functional connectivities ranging between 0.80 and 0.95. Values of the feed forward inhibition model parameter 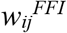 representing the strength of regional feedforward inhibitory input exhibited the most significant increase with age within the FP and DMN networks. A less pronounced, but significant, age-dependent increase of the inhibitory parameter values were seen in attention networks and no change within primary sensory networks.

**Conclusion:** Our study shows that high order (FP, DMN), attention and primary sensory networks exhibit distinct topographical and topological maturation trajectories. Moreover, brain network modeling revealed RSN-specific age-dependent inhibition trajectories, indicating that the model is able to reproduce and thus support candidate mechanisms of neurodevelopment.

**Graphical abstract:** Overview of the study combining empirical and modeling analyses of neuromaturation in 640 study participants from the HCP-D data set, age range 6-21 years old.

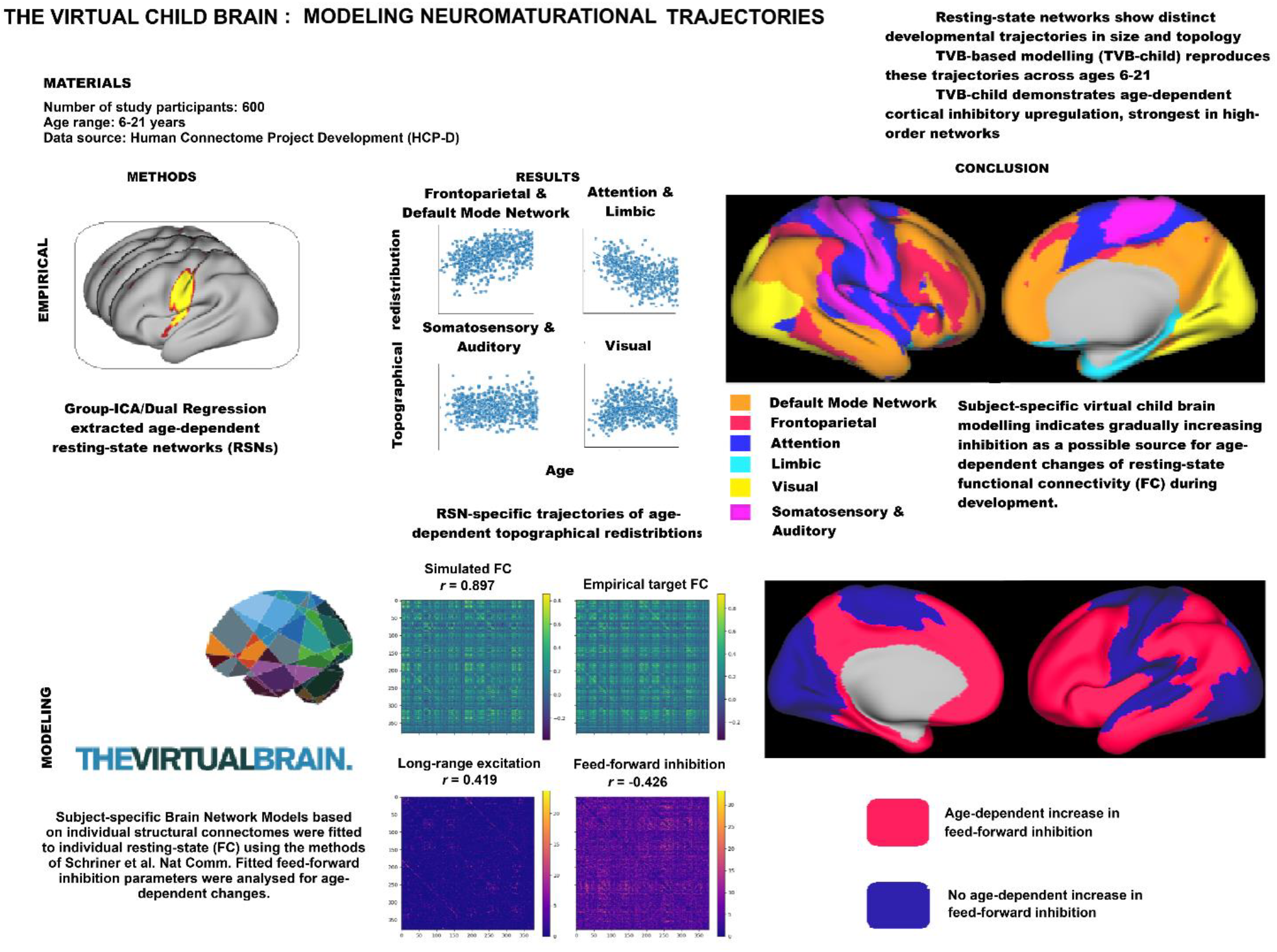

## Introduction

The first two decades of human life are characterized by remarkable neurodevelopmental changes towards the adult brain with its full cognitive capabilities (Konrad et al., 2013; Toga et al., 2006). Emergence of these capabilities occur along a genetically predetermined timeline with early childhood acquirement of primary sensory-motor abilities followed by a gradual development of executive functions until young adulthood (Giedd et al., 1999; Gogtay et al., 2004). Temporal developmental changes of spatial gradients, the so-called sensorimotor-association axis (SA-axis), ranging from brain regions linked to primary sensory brain functions, to brain regions linked to high-order functions have previously been characterized (Hill et al., 2010; Huntenburg et al., 2018). During development, postnatal myelination and cortical thinning are initiated in the posterior visual regions subsequently proceeding anteriorly to increasingly high-order brain regions (Gogtay et al., 2004; Huang et al., 2015; Lebel & Deoni, 2018). Correspondingly, childhood (6-12 years) resting-state network (RSN) activity is characterized by a strong primary sensory functional connectivity (FC). By late adolescence (12-18 years), the strongest FC connections become redistributed to association cortices (Dong et al., 2021). Childhood RSNs exhibit a high within-network connectivity with highly integrated local networks, subserving establishment of unimodal sensory and motor networks prior to adolescence onset and network refinement (Dong et al., 2021; Edde et al., 2021). Neurodevelopmental changes include a global decrease in both functional network modularity and local FC promoting global network integration. In addition, depending on RSN functionality, individual networks exhibit specific maturational trajectories. While resting-state FC between primary sensory networks increases, between-association cortices FC strength decreases with default mode (DMN) subnetworks becoming increasingly anti-correlated from the age 8 years until adulthood (Chai et al., 2014; Váša et al., 2020). These changes taking place until the third decade of life lead to an increased capability of association regions in mediating cognitive control and reasoning (Betzel et al., 2014; Fair et al., 2008; Luo et al., 2024; Marek et al., 2015; Sun et al., 2025; Wendelken et al., 2016). In summary, previous studies have characterized large-scale structural and functional changes in the SA-axis associated with neuromaturation and behavioral changes from childhood to adulthood. However, studying the underlying neurobiological mechanisms of these processes is difficult in humans. Rodent studies have demonstrated that gamma-aminobutyric acid (GABA) upregulation with subsequent inhibitory microcircuit maturation resulting in increased cortical inhibition levels constitute a hallmark of both developmental critical periods and experience-dependent synaptic pruning (Bosman et al., 2002; Erzurumlu & Gaspar, 2012; Runge et al., 2020). Non-human primate functional development is similarily linked to selective degeneration of excitatory synapses and sparing of inhibitory connections (Rakic & Burgeois, 1986). It can be hypothesized that similar neurobiological processes play a role in human neuromaturation unfolding along the SA-axis. A small number of experimental (Duncan et al., 2010; Fung et al., 2010) human studies have indicated a similar role for GABA upregulation in adolescent prefrontal development. Due to the limited possibilites to explore cortical inhibition and excitation *in vivo* in humans, computational modeling has been used to contribute to solving these questions (Larsen et al., 2022; Silveri et al., 2013). Computational brain network models generally have provided in the past important insights into principles of brain dynamics. The Virtual Brain (TVB) simulation platform enables modeling of whole-brain dynamics, linking large-scale brain dynamics to excitation-inhibition balance and have been widely used to model several neurological disorders including epilepsy and neurodegeneration (Ritter et al. 2013; Breakspear, 2017; Deco et al., 2014; Sanz-Leon et al., 2015; Schirner et al., 2023; Wong & Wang, 2006; Schirner et al, 2022; Koller et al 2024; Kashyap et al., 2025) making it a suitable tool for our study. By extending the computational modeling approach to neurodevelopment, our study adds a mechanistic framework for understanding how maturation shapes large-scale brain dynamics. Here we combined empirical data anylsis, i.e. group Independent Component Analysis/dual regression (gICA/DR) (Nickerson et al., 2017) for deriving age-dependent RSN FC and topological trajectories with subject specific large-scale dynamic computational brain modeling based on individual structural connectomes obtained from MRI data. The brain models were fitted to subject-specific functional connectivities obtained from functional magnetic resonance imaging (fMRI) data. The simulated FC reproduced observed empirical FC and the underlying alterations of model paramter values – representing regional inhibition and excitation levels - were analyzed to reveal possible causes of these observed developmental FC changes.

While the gICA/DR analysis demonstrated RSN-specific topographical and topological developemental trajectories, the simulations revealed that the value of inhibitory model paramters exhibited similar spatiotemporal trajectories, thus supporting the theory that altered inhibition levels play an important role in neuromaturaiton.

## Material and methods

### Ethics

A positive ethics vote for the present study has been obtained by the institutional Charité Ethics Committee (EA2/042/26).

### Resting-state network development

#### Material

FMRI and sMRI data from the lifespan Human Connectome Project Development (HCP-D) with 640 subjects, age range 6-21 years was utilized (Somerville et al., 2018). See Figure 1 for demographics. Participants were scanned with a customized Siemens 3 Tesla (T) Prisma scanner. The youngest subjects aged 6-7 years old were scanned with a pediatric 32-channel head coil. Both T1- and T2-weighted (T1w, T2w) sMRI scans were acquired with 0.8 millimeter (mm) isotropic voxels, Field of View (FOV) = 256 x 240 x 166 mm, matrix size = 320 x 300 x 208 slices and pixel bandwidth = 744 Hertz/Pixel (Hz/Px). T1w acquisition parameters include Repetition Time (TR)/Inversion Time (TI) = 2500/1000; Echo Time (TE) = 1.8/3.6/5.4/7.2 milliseconds (ms) and flip angle = 8 °. For T2w, TR/TE = 3200/564 ms and turbo factor = 314. Subjects older than 8 years old underwent 26 minutes of resting-state fMRI in four runs. Younger subjects underwent a reduced number of individual runs with total duration of 21 minutes of resting state fMRI. These were acquired with multiband, gradient recalled echo-planar imaging sequences. For details on data acquisition, see (Harms et al., 2018; Somerville et al., 2018). Both sMRI and fMRI data were pre-processed by the HCP consortium according to HCP minimal pre-processing guidelines. The pre-processed fMRI is organized in the HCP greyordinate CIFTI (Connectivity Informatics Technology Initiative) file system with cortical and subcortical surfaces represented as 91828 vertices (Glasser et al., 2013; Van Essen et al., 2012).

**Figure 1:**
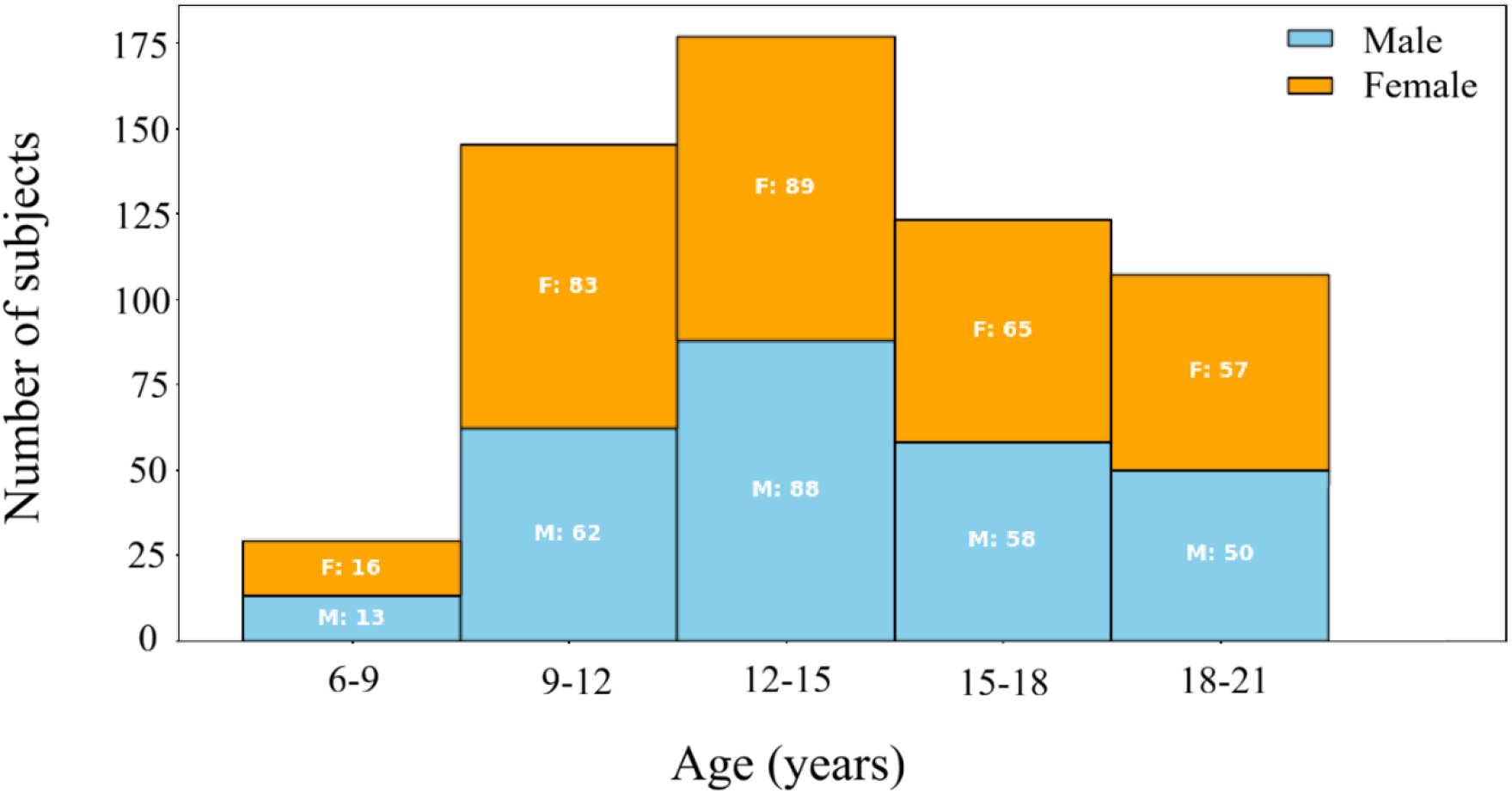
Demographic distribution: Histogram of gender and age study population distributions.

#### Group independent component analysis and Dual regression

Functional network topography is continuously refined through a gradual and individually variable neurodevelopment process (Cui et al., 2020). The resulting age-related spatial redistribution of large-scale networks cannot be captured by pre-defined, adult-derived brain parcellations (Bryce et al., 2021). To allow for characterization of such changes, gICA/DR was utilized to derive age-dependent trajectories of large-scale network spatial map distributions. The gICA components are computed by concatenation of all resting-state fMRI (rfMRI) time series with subsequent ICA computation. Hereby, a set of orthogonal spatial maps are derived, corresponding to population-specific resting-state networks (Nickerson et al., 2017). A two-step (dual) regression is subsequently performed. First, the gICA-derived, group-level spatial maps are used as spatial regressors in a general linear model (GLM) to estimate underlying time-series associated with the group-level ICs. These time-series are hereafter used as temporal regressors in the second GLM to compute subject-specific, IC-specific spatial maps. Group differences were inferred by comparison of the derived gICA/DR individual spatial maps (Reineberg et al., 2015). Here, gICA was computed on concatenated rfMRI time series of all subjects using FSL Multivariate Exploratory Linear Optimized Decomposition into Independent Components (FSL MELODIC) (Jenkinson et al., 2012). 30 gICs were computed. These gICs were used as input to HCPPipeline RSNregression.sh computing DR and thus outputting 30 gIC-DR spatial maps per subject. Z-score maps were computed using FSL MELODIC and thresholded at p<0.01 for all 30 group-level gICs as well as for all subject-specific gIC-DR spatial maps (Glasser et al., 2013). In conclusion, the gIC-DR computation results both in 30 gIC maps (thresholded at z-score > 1.96) common to the entire study population, and in 30x645 gIC-DR-maps, (thresholded at z-score > 1.96) i.e. individual spatial maps of the 30 gICs for each study participant.

#### Spatial and topological age-correlated RSN reorganization

GIC RSN affiliations were determined using spatial similarity analysis by computing cross-correlation using spatial Pearson correlation coefficient between each of the 30 gICs and the 7 Yeo atlas networks with the addition of an auditory network (Da Costa et al., 2011; Smith et al., 2009; Yeo et al., 2011). Dorsal and ventral attention networks were considered jointly as a common attention network. Each gIC was assigned to the network with which it exhibited the highest spatial Pearson correlation coefficient. Correlation coefficients < 0.25 were considered artefactual (Smith et al., 2009). For all analyses, gICs were averaged across the age-specific RSN to which they were assigned. To quantify any spatial redistribution of the RSNs, number of CIFTI vertices (Glasser et al., 2016) were averaged across the assigned gIC-DR maps for each subject. The number of CIFTI vertices per age-specific RSN per subject was then linearly regressed against age and in-scanner head motion as covariates:

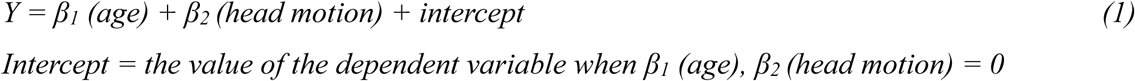

Variance defined as the difference between the observed outcome and the regression model’s prediction was used to determine whether data distributions were linear or non-linear. Data was considered linear if the residual was close to zero, and adding a non-linear term did not reduce the residual variance (Cook & Weisberg, 1995). In cases where the data distribution was non-linear, complementary non-linear regression was performed. A total of 21 regressions with 3 tests (topographical redistribution, degree, centrality) were performed on 7 RSNs with 2 covariates (age, head motion). 640 subjects were included in each test. All regression analyses were computed using Python library Scikit-learn (Pedregosa et al., 2011). Multiple comparisons were controlled using the Benjamini-Hochberg false discovery rate (BH-FDR) procedure to account for the number of tests across networks and metrics. This method adjusts p-values to yield q-values, controlling the expected proportion of false discoveries among significant findings. Statistical significance was defined as q < 0.05. To link age dependent functional changes to underlying structural properties, we analyzed changes in age-correlated cortical thinning. To this end, for each gIC, a cross-age (i.e. age independent) core region was defined as the set of CIFTI vertices with statistically significant BOLD correlation (z-score > 1.96) in gIC-DR maps across all ages. Correspondingly, gIC subregions that gained or lost statistically significant BOLD correlations (z-score > 1.96, corresponding to *p* < 0.05 (two-tailed) under the standard normal distribution) were defined as the developmental zone. See Figure 2 for a representative cross-age RSN with core and developmental zone. For all subjects and gICs, the average cortical thickness was computed for both the core and the developmental zone. Cortical thickness was extracted from the subject-specific HCP-D cortical thickness files (Glasser et al., 2013; Van Essen et al., 2012). The cortical thickness for both zones was averaged across the RSN and any age-dependent difference in cortical thickness trajectories between core and developmental zones were analyzed by comparing the rate of thinning between the zones using an interaction model equation:

**Figure 2:**
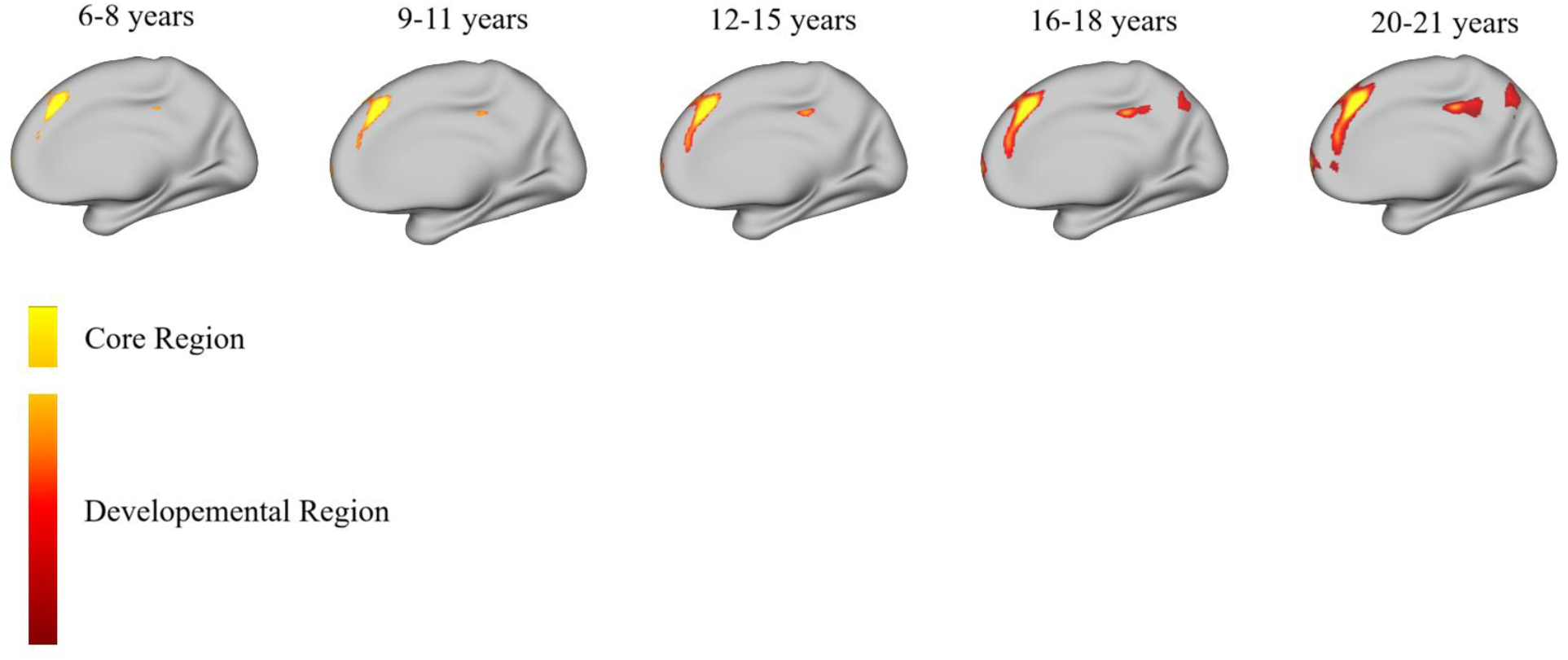
Representative gIC core and developmental regions: The core region is defined as the cortical area exhibiting a statistically significant BOLD correlations (z-score > 1.96) across all ages. The developmental region is defined as the cortical areas that gain, or lose, statistically significant BOLD correlations (z-score > 1.96) during development. Age-specific gIC2 presented as a representative case.

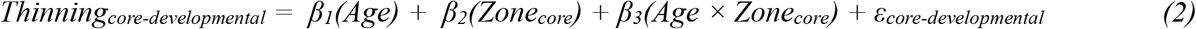

*β*_*1*_ : *Core zone thinning versus age slope*

*β*_*1*_ *+ β*_*3*_ *= Developmental zone thinning versus age slope*

*β*_*3*_ *= Interaction term reflecting difference in zone cortical thinning slope*

*ε*_*core-developmental*_ *= residual error reflecting the difference between the observed thickness and the model prediction* resulting in a total of 7 additional regression tests.

Within-network topological changes were quantified by computation of degree centrality and strength centrality averaged across all gICs within each age-specific RSN. Degree and strength centrality were computed by retaining top 5% of the strongest FC connections to construct a thresholded adjacency matrix. Degree centrality was defined as the number of suprathreshold connections for each vertex. Strength centrality was defined as the sum of the suprathreshold connections (Van Den Heuvel & Sporns, 2013)

For a network *G = (V, E)*, |*V*| *=* number of CIFTI vertices, |*E*| = number of edges, degree centrality *C*_*D*_ of an individual vertex *v* is defined as

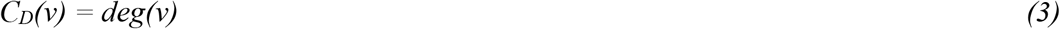

Strength centrality *C*_*S*_ for a CIFTI vertex *v* with adjacency matrix *W = (w*_*ij*_) is defined as

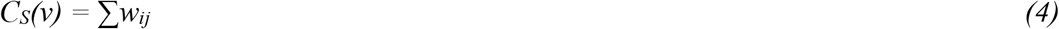

Degree centrality and strength centrality was averaged across each age-specific RSN for all subjects. Regression of age-specific RSN degree and strength centrality against age was computed according to

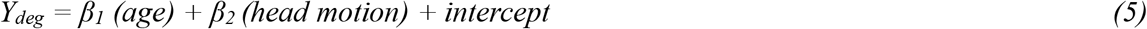

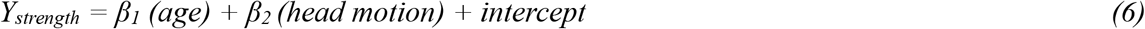

In addition to analyzing within-network topological changes, age-related changes in between-network connectivity were characterized. One-to-all gIC FC strength was computed as the mean Pearson correlation between one gIC, and all other gICs (Lou et al., 2023). This was done for all gICs and for all subjects. Regression analysis against age was computed for each gIC.

### Virtual Child Brain Network Model

#### Diffusion weighted data and generation of structural connectomes

For details on sMRI and fMRI data processing, see section <u>Resting-state network development.</u>

#### Material

Diffusion-weighted data (DWI) from the HCP-D project was for all study subjects acquired with 185 directions on 2 shells with b=1500 and 3000 s/mm^2^ within the HCP-D consortium (Harms et al., 2018). DWI data was pre-processed by the authors in accordance with the HCP Diffusion Preprocessing pipeline. B0 values were normalized across all runs followed by subject motion artifact removal and echo-planar imaging (EPI), eddy current distortion and gradient non-linearity corrections (Glasser et al., 2013; Van Essen et al., 2012). Whole-brain tractography was computed using MRTrix software. Pre-processed DWI data was co-registered with sMRI data followed by computation of spherical deconvolution-based fiber-orientation distributions (Jeurissen et al., 2014). Hereafter, the whole-brain tractography was parcellated according to the Glasser atlas to create a structural connectome (Glasser et al., 2016).

#### Brain Network Model

TVB simulates large-scale brain networks by coupling mesoscopic neural mass models modeling regional brain activity using subject-specific tractography-derived structural connectome data to weight inter-nodal edges (Ritter et al. 2013; Sanz-Leon et al. 2013; Sanz-Leon et al., 2015; Schirner et al. 2022). The TVB-Child model aims to explore the role for regional cortical inhibition in neurodevelopment. To this end, a previously developed TVB model with long range excitation and feed forward inhibition was utilized (Schirner et al., 2023). A dynamical mean field reduction of a local spiking network with leaky integrate-and-fire neurons simulating coupled excitatory and inhibitory populations extended by two parameters reflecting long-range excitation 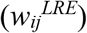 and feedforward inhibition 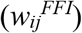 was utilized to simulate local brain activity (Deco et al., 2014; Schirner et al., 2023). These two additional parameters were tuned individually for each node, resulting in regional cortical inhibition levels. The brain network model was defined as follows (Schirner et al. 2023)

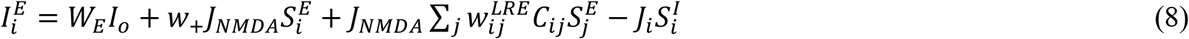

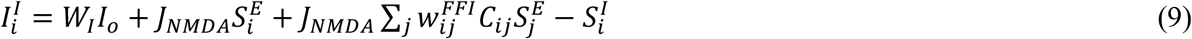

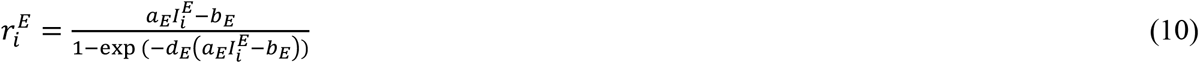

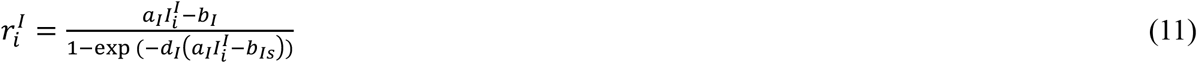

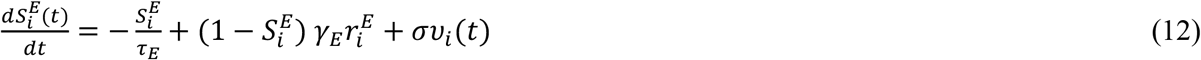

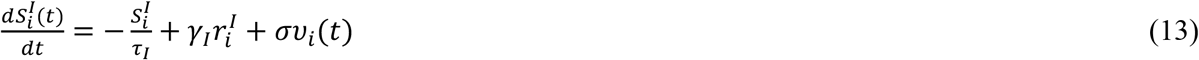

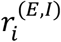=Excitatory (*E*) and inhibitory (*I*) population firing rate of cortical region *i*. 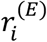 is set to 4 Hz.

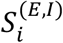=Average excitatory and inhibitory synaptic gating of region *i*.

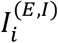=Region *i* input current

*W*_(*E,I*)_*I*_*o*_ =External input currents to excitatory and inhibitory populations

*w*_+_ =Local excitatory recurrence

*J*_(*NMDA,i*)_=Excitatory and local feedback inhibitory synaptic coupling, respectively

*C*_*ij*_ =SC with shape (number of regions × number of regions)

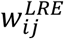=Long-range excitation with shape as SC

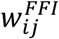=Feed-forward inhibition with shape as SC

*τ*_(*E,I*)_, *γ*_(*E,I*)_=Time and rate of saturations of excitatory and inhibitory activity.

*v*_*i*_(*t*)= Noise from standard normal distribution.

One TVB-Child model was set up for each study subject. Network edges of each such model were weighted by the individual subject’s structural connectivity, resulting in a model informed by age-specific white matter properties. The resulting excitatory synaptic activity 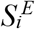 used as input to a Ballon-Windkessel model describing the relationship between neural activity and the BOLD fMRI signal. Hereafter, the model outputs one simulated fMRI signal per Glasser atlas parcel. (Deco et al., 2014; Glasser et al., 2016; Jeurissen et al., 2014).

#### Generation of empirical and simulated functional connectivities

An empirical FC for each subject was constructed by parcellation of the fMRI time series using the Glasser atlas (Glasser et al., 2016). Here after, Pearson correlations between all parcels were computed. To this aim, HCP pipeline -cifti-parcellated followed by -cifti-correlation was used (Glasser et al., 2013). The simulated FC was constructed by computation of Pearson correlations between all simulated fMRI signals.

#### Model fitting

The resulting simulated fMRI FC was fitted to the corresponding subject-specific fMRI FC by region-wise tuning of the long-range-excitation versus feed-forward-inhibition ratios using the algorithm described in (Schirner et al. 2023). Thus, the TVB-Child model informed by subject-specific white matter information, tuned regional inhibition levels to output subject-specific fMRI FC. Following Schirner et al. 2023, Feedback Inhibition Control (FIC) was used to regulate synaptic firing rates (Deco et al., 2014; Schirner et al., 2023). Gradient descent was used to tune parameters (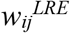 and 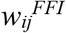) to achieve a high correlation between the empirical and simulated FCs. The model parameters (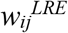 and 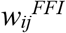) were fitted with 300 iterations and simulation length 60 000 seconds to achieve stable empirical-simulation FC correlations. After each simulation run, global model performance was evaluated using entry-wise root-mean square (RMSE) between the empirical target FC and the simulated FC. Parameters were hereafter updated using a learning rate = 0.2. The fitting algorithm was implemented in Python with a differentiable backend based on JAX and PyTensor libraries (Frostig et al., 2018). Thus, through simulation, we inferred the region-pair-wise long-range-excitation and feed-forward-inhibition ratios (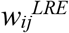 and 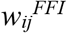) leading to the best fit of simulated and empirical subject specific FCs. Repeating the fitting algorithm with different initializations did not yield converging parameters; however, we evaluated the robustness of the fitted models by running multiple simulations with identical noise levels. Across 1000 repetitions, the predicted fMRI time series were highly consistent (pairwise correlations > 0.99), indicating that the model outputs are stable despite parameter variability. The fitting and validation procedure followed (Schirner et al., 2023).

#### Modeling of age-correlated trajectories of cortical inhibition

To analyze a possible statistical relationship between the TVB-Child model inferred parameters (representing the ratios of long-range-excitation versus feedforward-inhibition) and the gIC/DR cross-age RSN features, the Glasser atlas parcels were assigned to one of Yeo 7 atlas regions frontoparietal (FP), default mode (DMN), attention (ventral and dorsal attention), somatosensory (SM), visual or limbic networks with the added auditory network (Da Costa et al., 2011; Yeo et al., 2011). Network assignment between Glasser and Yeo 7 atlas was done by computing HCP surface CIFTI vertex overlap (Glasser et al., 2016; Yeo et al., 2011) in Glasser parcellation and Yeo 7 atlas 32k_fs_LR.dlabel files. These are surface-based parcellation maps in CIFTI format that assign each vertex on the 32k-resolution left-right symmetric cortical mesh to a specific anatomical or functional label. For each Glasser parcel, the number of vertices belonging to each parcel was quantified. Here after, the number of such vertices belonging to each Yeo 7 network was computed and the individual Glasser parcel was assigned to the Yeo 7 network containing the largest proportion of vertices. For all age-specific RSNs and for all subjects, within-network cortical inhibition levels of network *k* were computed as

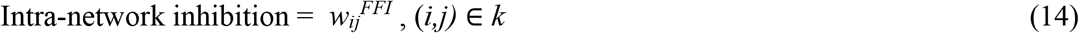

Similarly, between-network cortical inhibition levels between networks *k, l* was computed as

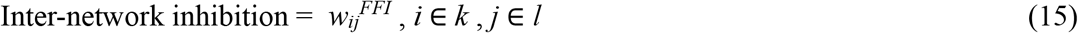

Correlations between age and cortical inhibition was estimated using linear regression as described in *Spatial and topological age-correlated RSN reorganization*.

## Results

### Resting-state network development

Cross-age RSN development was estimated using a combined gICA/DR approach. GICA based upon a concatenation of all rfMRI time series was used to create individual spatiotemporal IC maps (gICs) from which age-related changes in cross-age RSN topography and topology were inferred by linear and non-linear regression analysis. One gIC (gIC 14) did not exhibit any significant BOLD correlation z-score and was excluded from further analysis. One gIC (gIC 25) overlapped exclusively with the primary auditory region (da Cost a) and was thus classified as auditory network. The remaining 28 gICs were classified as FP, DMN, attention (both dorsal and ventral attention), limbic, visual and SM RSNs by computation of spatial Pearson correlation coefficients between individual gICs and the Yeo 7 RSN atlas (Yeo et al., 2011). Of these 28 gICs, eleven correlated strongest with high-order association networks FP (4 gICs) and DMN (7 gICs); five with an attention network and one with the limbic network; eleven with primary sensory networks, that is 5 gICs with the SM RSN and 6 gICs with the visual RSN. Similar RSN subdivisions in adults have previously been reported using gICA/DR (Reineberg et al., 2015). GIC RSN subregions are displayed in Figure 3 with all gIC components grouped according to network affiliation.

**Figure 3:**
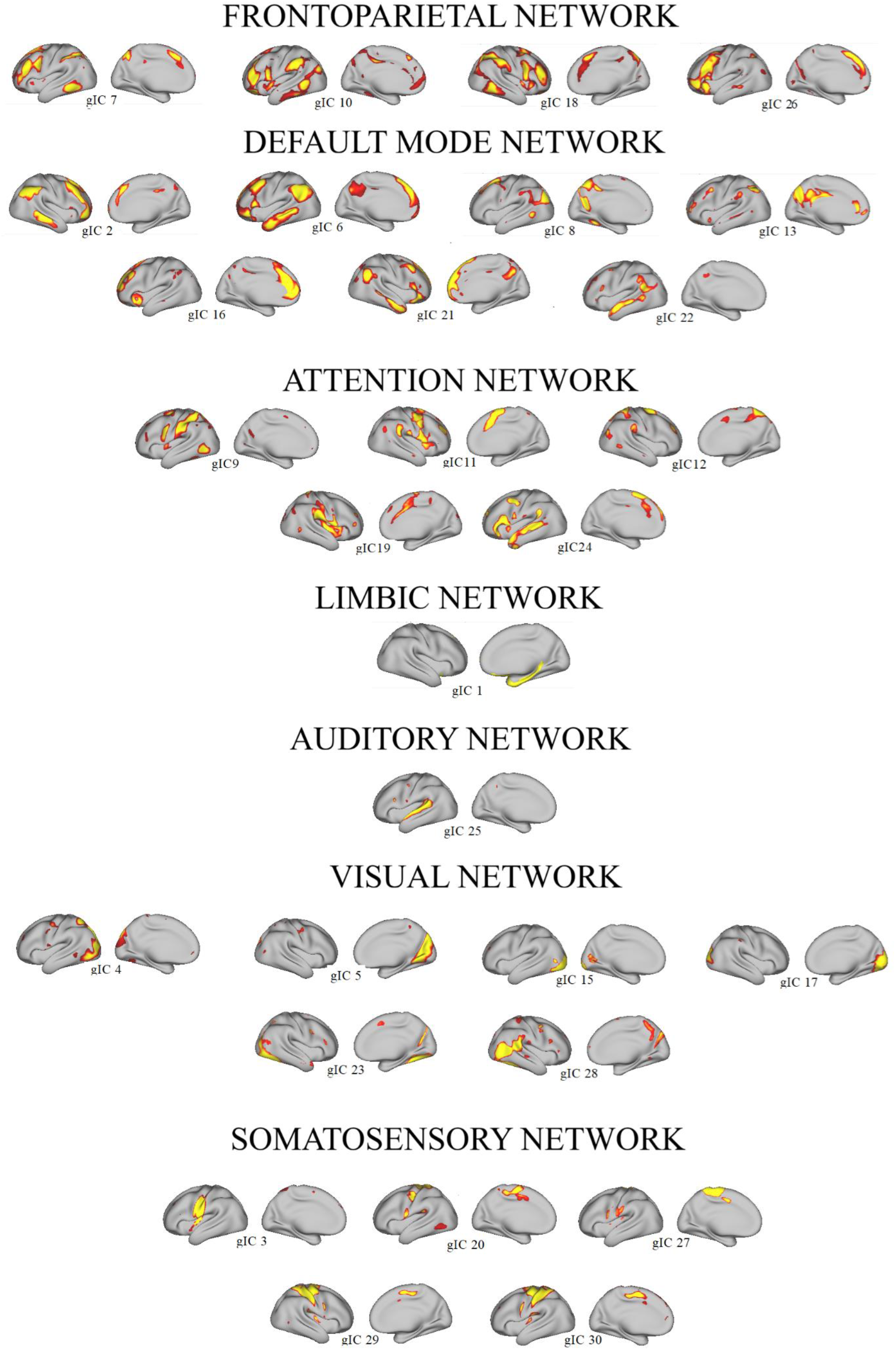
Spatial distribution of group-common independent components: Statistically significant (z-score > 1.96) group-common independent components group by affiliation to resting state networks (RSN) frontoparietal, default mode, attention (including both ventral and dorsal attention), limbic, visual and somatosensory networks according to Pearson correlation with the Yeo (Yeo et al., 2011) 7 RSN atlas. In addition, gIC25 was classified as auditory network due to overlap with primary auditory cortex. gIC2 and gIC6 exhibited only uni-hemispheric temporal correlations. Remaining gICs exhibited symmetric bi-hemispheric temporal correlations; the hemisphere exhibiting the most prominent temporal correlation pattern is presented here. GIC14 with no significant BOLD temporal correlations was not included.

### Within-network connectivity

To characterize cross-age RSN spatial reorganization, the number of vertices on the cortical surface, averaged across all gICs within the respective age-specific RSN, exhibiting a statistically significant z-score were regressed against age and in-scanner head motion. No significant correlations were found between head motion and significant z-score values. Regression against age demonstrated that association cross-age RSNs FP and DMN exhibit a linear increase (regression p-value < 0.05; BH-FDR q-value < 0.05) in number of vertices with a significant BOLD z-score. Attention and limbic networks exhibited the opposite trend with an age dependent linear decrease of the number of vertices with a significant BOLD z-score (regression p-value < 0.05; BH-FDR q-value < 0.05). This shift in network distribution reflects an age-dependent spatial increase in cortical areas (Harris et al., 2011) subserving association networks, and a corresponding spatial decrease in attention and limbic network cortical representation. Primary sensory SM and auditory networks did not exhibit any significant spatial redistribution while the visual network exhibited an inverted U-shaped (regression p-value < 0.05) in number of vertices with significant SM cross-age RSN BOLD z-score. In conclusion, cortical representation of association networks expanded while attention networks underwent age-dependent refinement. Except for visual networks, primary sensory networks remained stable. See Figure 4 for scatter plots with significant (p-value < 0.05) regression trajectories, correlation coefficients and coefficients of determinations of the number of vertices with significant BOLD correlations (z-score > 1.96) across age as well as cortical maps of the anatomical distribution of these findings. See Table 1 for details on regression coefficients, p-values and BH-FDR q-values for topographical redistribution across age. To link functional maturation to any underlying structural maturation, we analyzed cortical thickness changes in cross-age RSN subregions that remained stable (core region) compared to cross-age RSN regions that gained or lost cross-age RSN specific BOLD correlations (developmental region) with age (see Figure 2 for representative core and developmental regions). Cortical thinning is a well described feature of human neuromaturation (Giedd et al., 1999). Within cross-age RSNs that grew with age, cortical thinning was most pronounced within the developmental region (DMN regression coefficient β = −4.7e-3; FP regression coefficient β = −4.3e-3; all p-values < 0.05) compared to core region (DMN regression coefficient β = −1.1e-3; FP regression coefficient β = −1.2e-3; all p-values < 0.05). In conclusion, functional and structural remodulation were coupled within high-order cross-age RSNs. Within cross-age RSNs that diminished with age, cortical thinning was most pronounced within the core region (attention regression coefficient β = −2.7e-3; limbic regression coefficient β = −2.8e-3; all p-values < 0.05) compared with the developmental region (attention regression coefficient β = −1.1e-3; limbic regression coefficient β = −1.2e-3; all p-values < 0.05). Thus, the cross-age RSN subregions that survived network refinement were characterized by more prominent structural remodulation than the cross-age RSN subregions that underwent network pruning. See Figure 4, panel (ii).

**Table 1.**
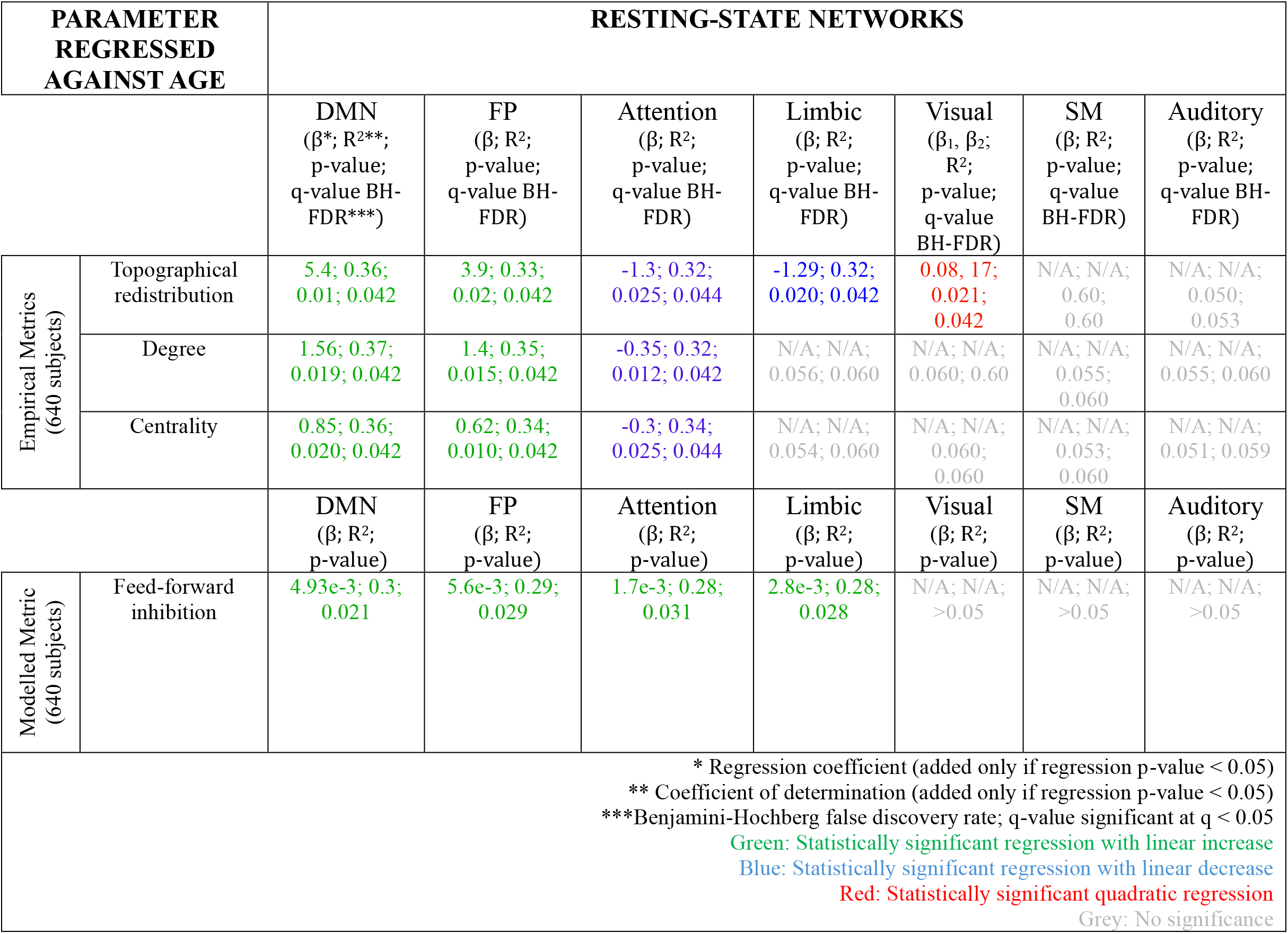
Regression coefficients, p-values and BH-FDR q-values for topographical redistribution across age.

| PARAMETER REGRESSED AGAINST AGE |  | RESTING-STATE NETWORKS |  |  |  |  |  |  |
| --- | --- | --- | --- | --- | --- | --- | --- | --- |
| | | DMN<br>($\beta^*$; $R^{2**}$;<br>p-value;<br>q-value BH-FDR***) | FP<br>($\beta$; $R^2$;<br>p-value;<br>q-value BH-FDR) | Attention<br>($\beta$; $R^2$;<br>p-value;<br>q-value BH-FDR) | Limbic<br>($\beta$; $R^2$;<br>p-value;<br>q-value BH-FDR) | Visual<br>($\beta_1, \beta_2$;<br>$R^2$;<br>p-value;<br>q-value BH-FDR) | SM<br>($\beta$; $R^2$;<br>p-value;<br>q-value BH-FDR) | Auditory<br>($\beta$; $R^2$;<br>p-value;<br>q-value BH-FDR) |
| Empirical Metrics<br>(640 subjects) | Topographical redistribution | 5.4; 0.36;<br>0.01; 0.042 | 3.9; 0.33;<br>0.02; 0.042 | -1.3; 0.32;<br>0.025; 0.044 | -1.29; 0.32;<br>0.020; 0.042 | 0.08, 17;<br>0.021;<br>0.042 | N/A; N/A;<br>0.60;<br>0.60 | N/A; N/A;<br>0.050;<br>0.053 |
|  | Degree | 1.56; 0.37;<br>0.019; 0.042 | 1.4; 0.35;<br>0.015; 0.042 | -0.35; 0.32;<br>0.012; 0.042 | N/A; N/A;<br>0.056; 0.060 | N/A; N/A;<br>0.060; 0.60 | N/A; N/A;<br>0.055;<br>0.060 | N/A; N/A;<br>0.055; 0.060 |
|  | Centrality | 0.85; 0.36;<br>0.020; 0.042 | 0.62; 0.34;<br>0.010; 0.042 | -0.3; 0.34;<br>0.025; 0.044 | N/A; N/A;<br>0.054; 0.060 | N/A; N/A;<br>0.060;<br>0.060 | N/A; N/A;<br>0.053;<br>0.060 | N/A; N/A;<br>0.051; 0.059 |
| Modelled Metric<br>(640 subjects) | | DMN<br>($\beta$; $R^2$;<br>p-value) | FP<br>($\beta$; $R^2$;<br>p-value) | Attention<br>($\beta$; $R^2$;<br>p-value) | Limbic<br>($\beta$; $R^2$;<br>p-value) | Visual<br>($\beta$; $R^2$;<br>p-value) | SM<br>($\beta$; $R^2$;<br>p-value) | Auditory<br>($\beta$; $R^2$;<br>p-value) |
|  | Feed-forward inhibition | 4.93e-3; 0.3;<br>0.021 | 5.6e-3; 0.29;<br>0.029 | 1.7e-3; 0.28;<br>0.031 | 2.8e-3; 0.28;<br>0.028 | N/A; N/A;<br>>0.05 | N/A; N/A;<br>>0.05 | N/A; N/A;<br>>0.05 |
| * Regression coefficient (added only if regression p-value < 0.05)<br>** Coefficient of determination (added only if regression p-value < 0.05)<br>***Benjamini-Hochberg false discovery rate; q-value significant at $q < 0.05$<br>Green: Statistically significant regression with linear increase<br>Blue: Statistically significant regression with linear decrease<br>Red: Statistically significant quadratic regression<br>Grey: No significance |  |  |  |  |  |  |  |  |

**Figure 4:**
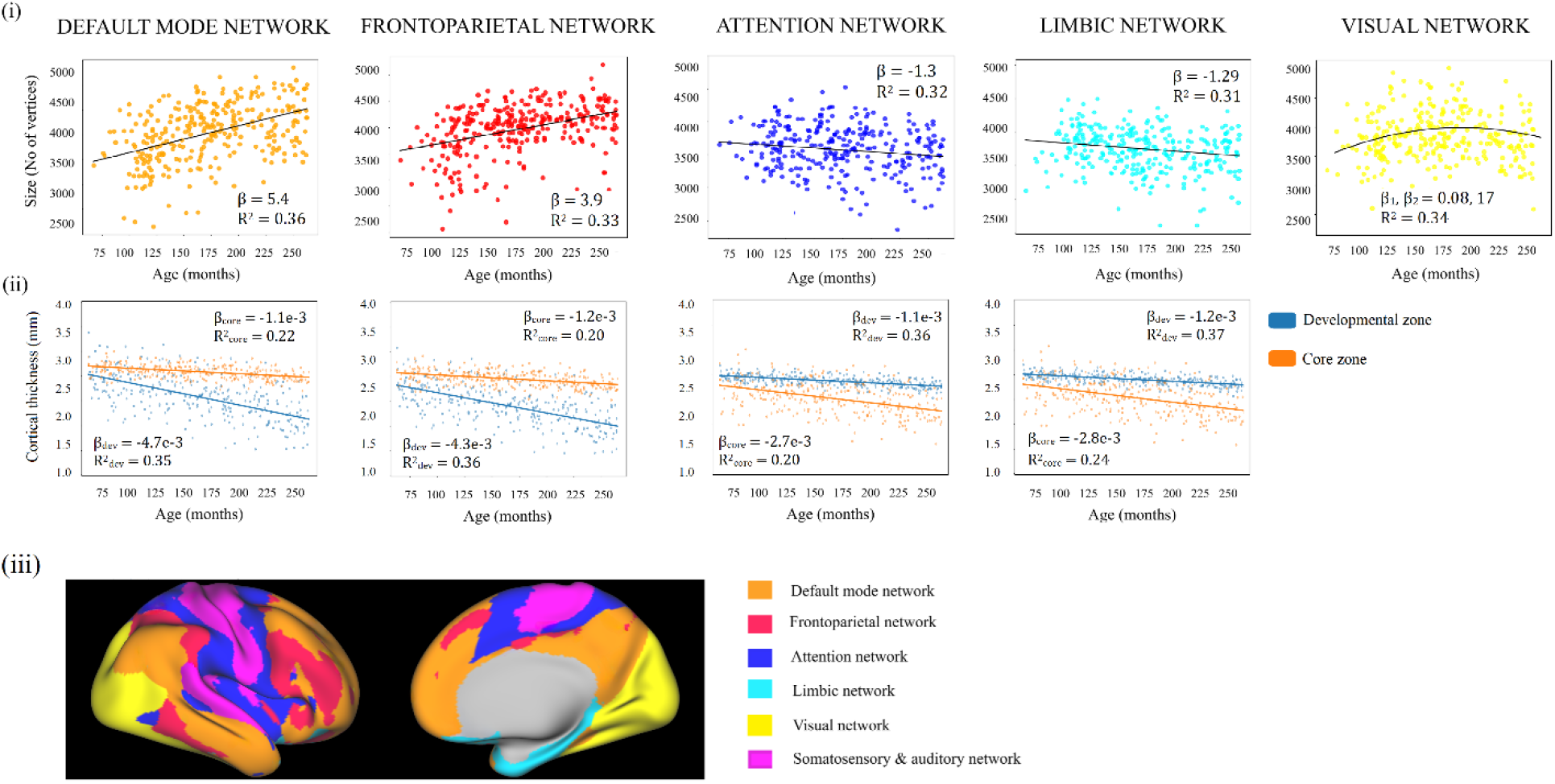
Maturational trajectories of spatial redistribution. **(i)** Individual data points represent subject group Independent/Component/Dual Regression (gIC/DR) map (threshold z-score > 1.96) number of vertices with RSN specific significant resting state BOLD correlations averaged across gICs within Default Mode Network (DMN), Frontoparietal network (FP), Attention network, Limbic and Visual Networks, respectively regressed against age (unit: months). Statistically significant linear (p-value < 0.05) regression trajectories are presented with correlation coefficient β and determination of coefficient of determination *R*^*2*^. Statistically significant (p-value < 0.05) quadratic regressions are presented with quadratic and linear coefficients (β_1_, β_2_). RSNs (auditory and somatosensory networks) with non-significant changes in BOLD response were not presented. (ii): Cortical thinning (millimeter, mm) versus age for developmental and core regions. (iii): Anatomical distributions of the analyzed RSNs.

To characterize cross-age RSN topological changes, cross-age RSN degree and strength centrality were analyzed. High-order cross-age RSNs FP and DMN exhibited an increase in both degree and strength centrality (regression p-value < 0.05). The cortical representation expansion was thus parallelled by forming network hubs with increased information integration across larger cortical areas. Alongside network size pruning, attention cross-age RSNs exhibited a decrease (regression p-value < 0.05) in both degree and strength centrality. See Figure 5 and Figure 6 for cross-age RSN scatter plots and trajectories for all statistically significant regressions, as well as inflated surfaces demonstrating the anatomical distribution of topological changes. See Table 1 for details on regression coefficients, p-values and BH-FDR q-values for topographical redistribution across age.

**Figure 5:**
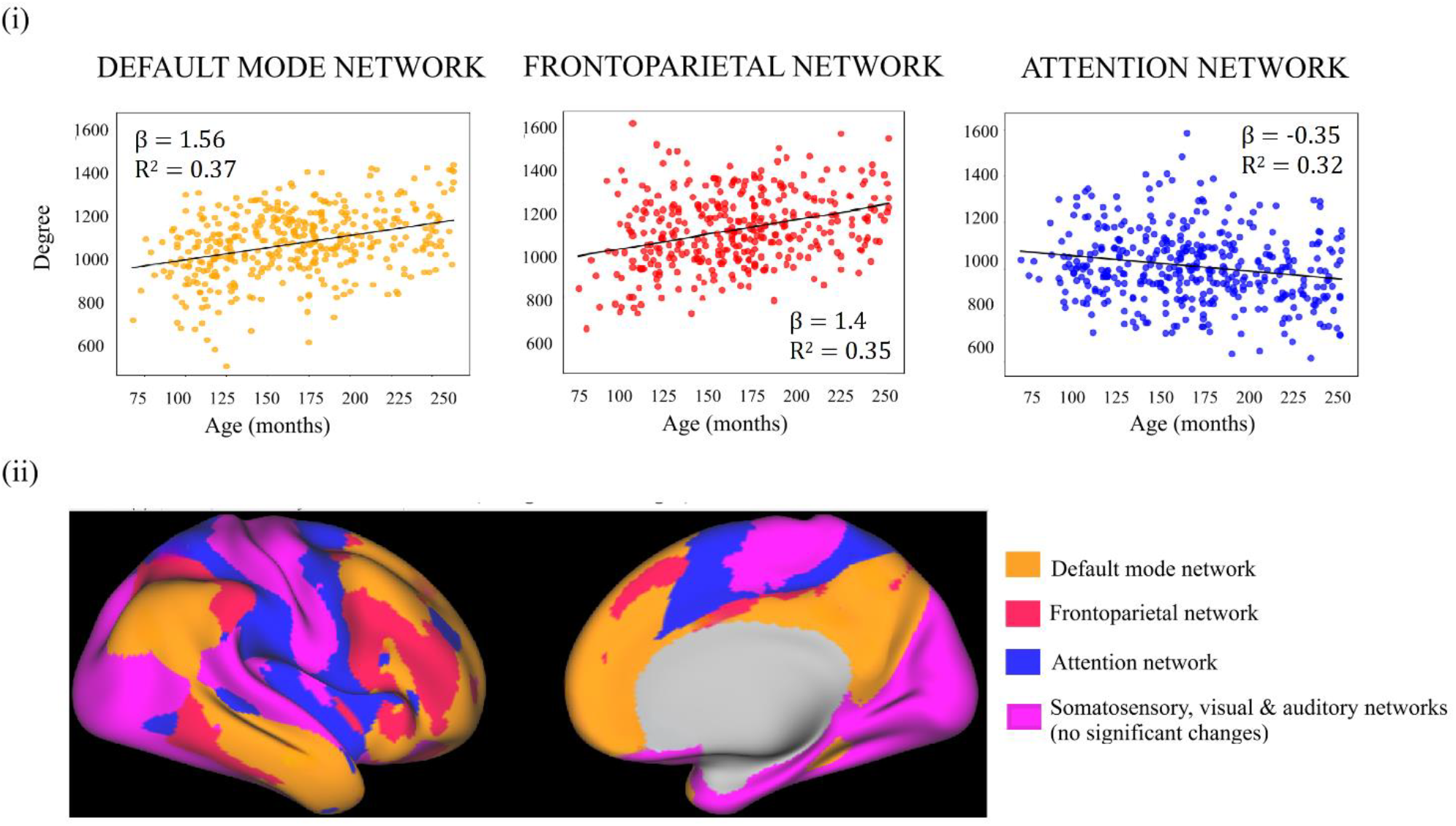
Maturational trajectories of degree centrality. **(i):** Individual data points correspond to single-subject average vertex degree centrality averaged across gICs belonging to respective RSNs regressed against age (unit: months). Only statistically significant (p-value < 0.05) are presented. Regression trajectories are presented with correlation coefficient β and determination of coefficient of determination *R*^*2*^. **(ii):** Anatomical distributions of the analyzed RSNs.

**Figure 6:**
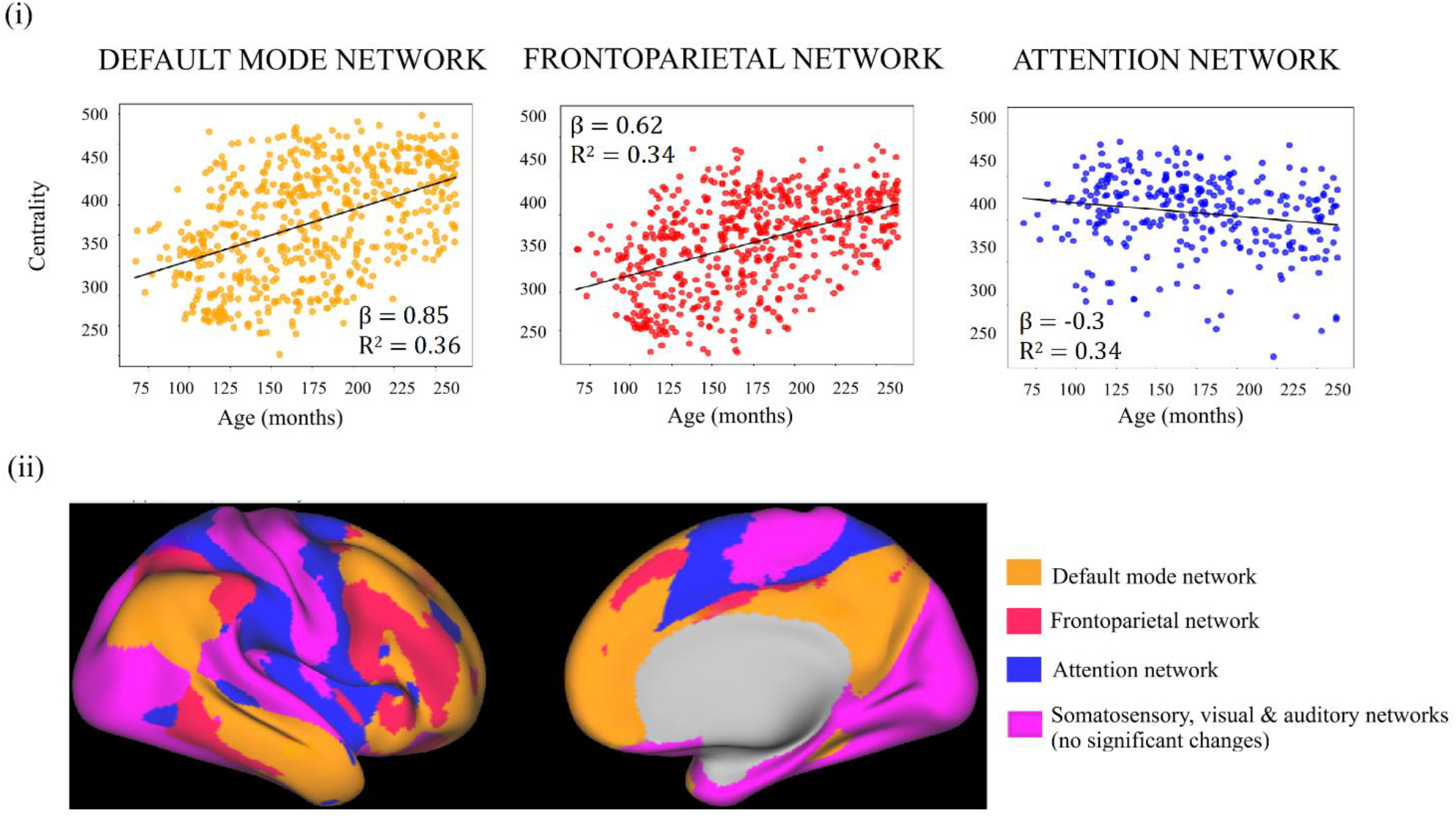
Maturational trajectories of strength centrality. **(i):** Individual data points correspond to average vertex-wise strength centrality averaged across gICs belonging to respective RSNs regressed against age (unit: months). Only statistically significant (p-value < 0.05) are presented. Regression trajectories are presented with correlation coefficient β and determination of coefficient of determination *R*^*2*.^. **(ii):** Anatomical distributions of the analyzed RSNs.

In summary, the results suggest cross-age RSN-specific topographical and topological redistribution, with high-order FP and DMN networks expanding and becoming increasingly important in information integration. Attention network underwent pruning, while primary sensory networks, except for visual networks exhibiting a U-shaped trajectory, remained stable.

### Between-network connectivity

Between-network topology and organization is known to shift during development to subserve cognitive maturation (Song et al., 2024). In order to analyze whether these changes followed RSN-specific trajectories, we charcterized between-gIC connectivity. Quantifying FC strength between individual association gICs and all other individual gICs demonstrate that the regression coefficients (p-value < 0.05) of inter-gIC FC across age exhibit range [−5.5 – 1.3], median −1.2, mean −2. FC strength correlation coefficients (p-value < 0.05) between attention gICs and attention and primary gICs exhibit range [−1.3 – 1.2], median −0.3 and mean −0.3. FC strength regression coefficients (p-value < 0.05) between primary gICs have range [−1.2 – 0.3], median 0.1 and mean 0.1. Thus, high-order RSN between-network connectivity decreased most prominently with age, indicating a maturational network segregation. Connectivities between attention and primary sensory networks exhibited a mean increase in FC strength, consistent with previous findings (Luo et al., 2024). See Figure 6 for all between-gIC changes in FC strength regression coefficients against age.

### Virtual Child Brain network model

The gICA/DR showed cross-age RSN-specific topographical and topological redistributions consistent with the existing literature (Hill et al., 2010; Huntenburg et al., 2018; Luo et al., 2024).

With computational brain network modeling we aimed to reproduce these alterations of FC during age dependent neuromaturation. Subject-specific TVB-Child models were fitted to individual study participants’ FCs, resulting in simulated fMRI FC matrices highly correlated to the corresponding empirical fMRI FC (correlation coefficient *r =* [0.85-0.95]; see Figure 7, panel (i) for a histogram of all study participant correlation coefficients. Global feed-forward inhibition averaged across all simulated regions increased (regression coefficient 2e-3) while global long-range excitation averaged across all regions decreased (regression coefficient −1.5e-3); see Figure 7, panel (ii)-(iii).

**Figure 7:**
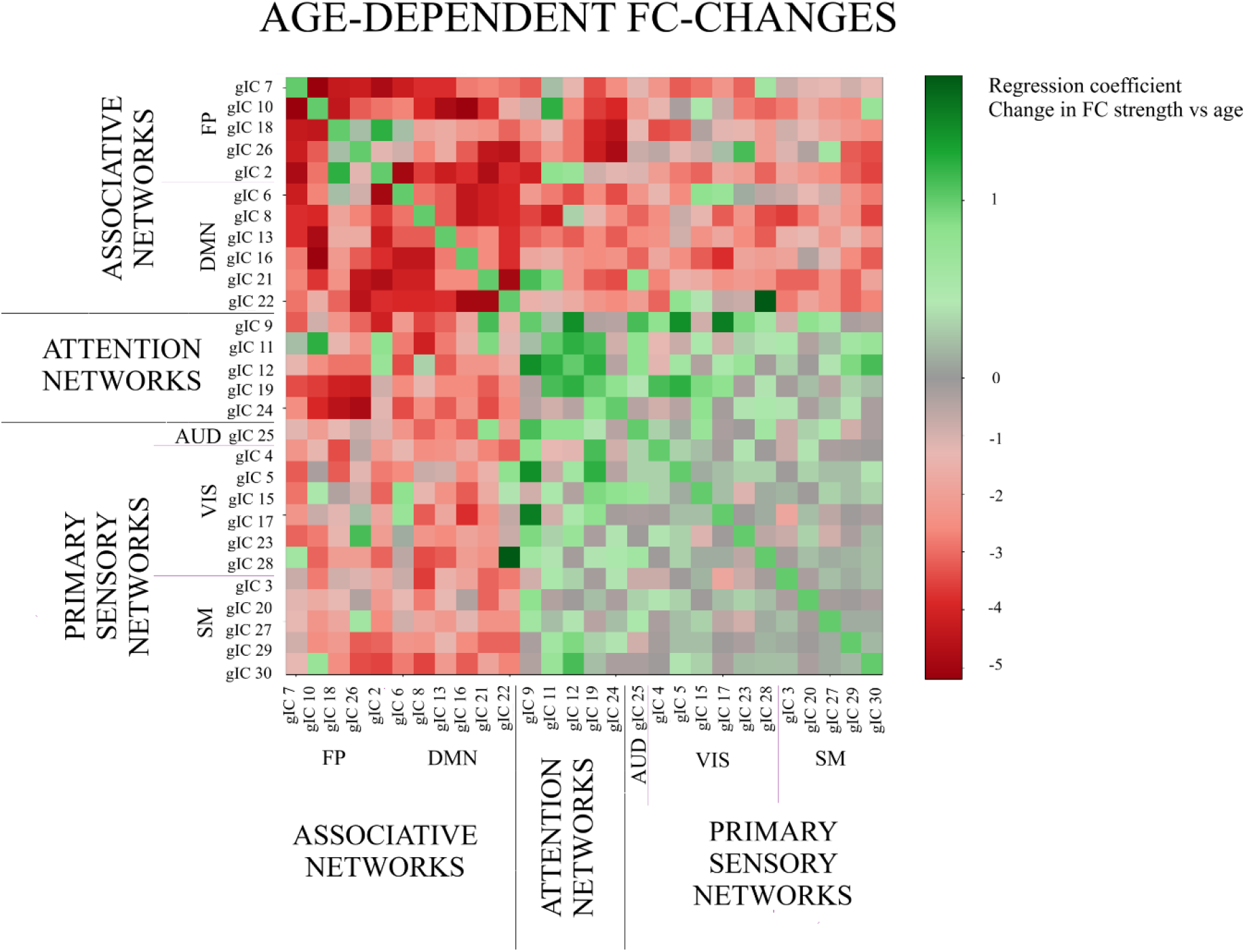
Age dependent functional connectivity strength changes: Matrix entry (*j,i*) represents the regression coefficient of FC strength vs age between gIC *i* and gIC *j*.

### Within- and between network inhibition

After fitting individual FC matrices, we explored whether the empirically observed relationship between FC and SA-axis position was paralleled by altered model inhibition parameters across age. Since previous studies have shown that GABA upregulation mediates neurodevelopment and activity-depended plasticity (Bosman et al., 2002; Erzurumlu & Gaspar, 2012; Runge et al., 2020), we analyzed spatiotemporal gradients of the region-wise fitted long-range excitation and feed-forward inhibition parameters. To compare model output with the gICA/DR results, the Glasser parcellation of the simulated data was subdivided into FP, DMN, attention, limbic, SM, auditory and visual networks. We found that association FP and DMN, attention and limbic networks exhibited a significant increase in within-network inhibition levels with age (regression coefficients β = 5.6e-3 – 1.7e-3, R^2^ = 0.28-0.3, p-value < 0.05). The most prominent changes were seen within association networks. Primary sensory networks did not exhibit any significant changes in within-network inhibition levels. See Figure 8 for all scatter plots, significant trajectories and the anatomical distribution of age-dependent inhibitory upregulation.

**Figure 8:**
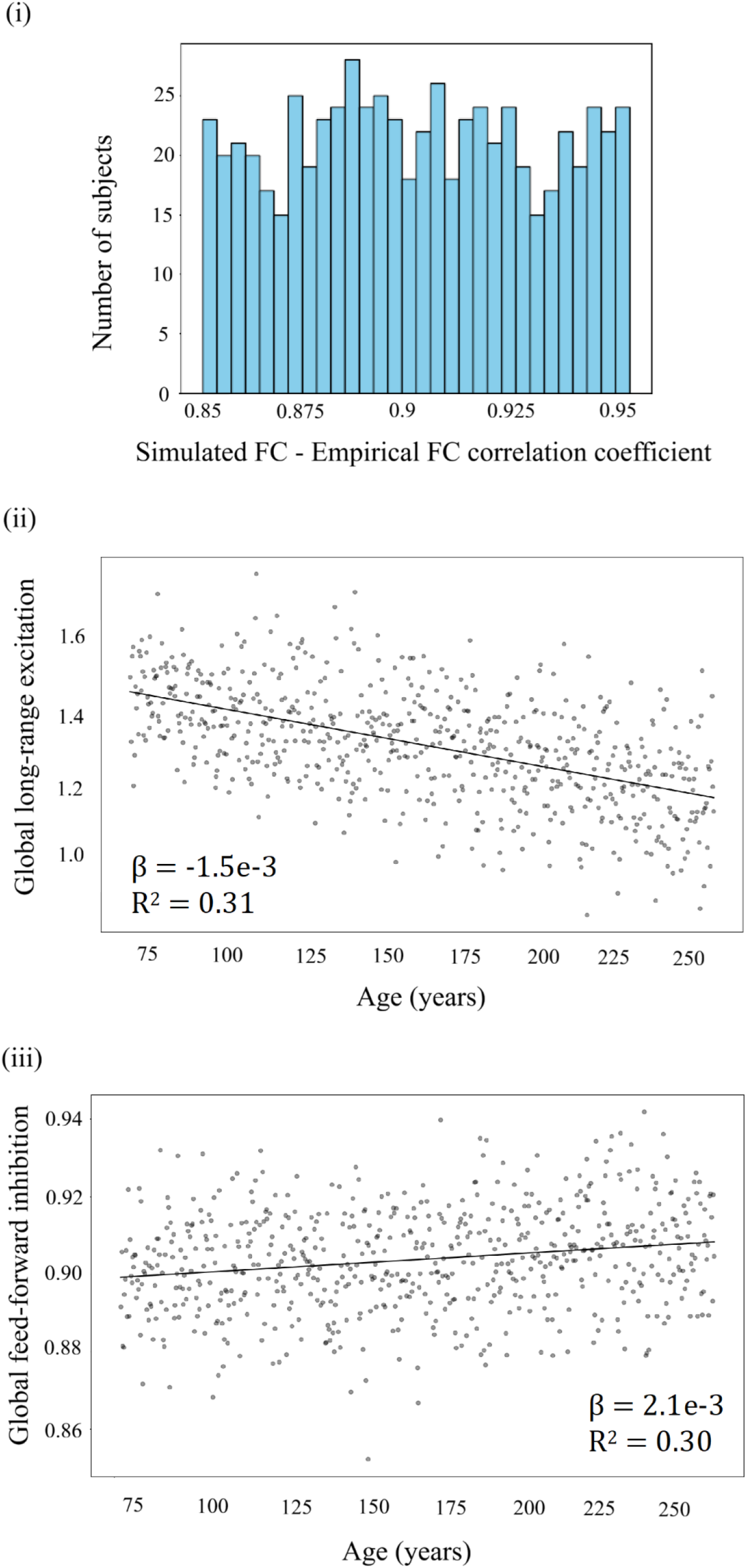
Simulation parameters. **(i):** Histogram of all subject simulated-empirical functional connectivity correlation coefficients (range 0.85-0.95). **(ii):** Individual data points represent subject-specific, global long-range excitation parameters *w*^*LRE*^ averaged across all simulated Glasser parcels. Regression coefficient against age *β* =-1.5e-3; coefficient of determination R^2^ =0.31 (p-value < 0.05). **(iii):** Individual data points represent subject-specific, global feed-forward parameters *w*^*FFI*^ averaged across all simulated Glasser parcels. Regression coefficient against age *β* =2.1e-3; coefficient of determination R^2^ =0.30 (p-value < 0.05).

Between-network inhibition levels regressed against age demonstrated that connections between FP and all other networks, and between DMN and all non-primary sensory networks were marked by the most pronounced inhibitory upregulation (regression coefficients β = [6e-3 – 8e-3], p-value < 0.05). Between-network connections involving attention or limbic networks were characterized by a less prominent increase in inhibition (regression coefficients β = [2e-3 – 5e-3], p-value < 0.05). There were no significant changes in inhibition between primary sensory networks. See Figure 9 for all simulated between-network inhibition levels.

**Figure 9:**
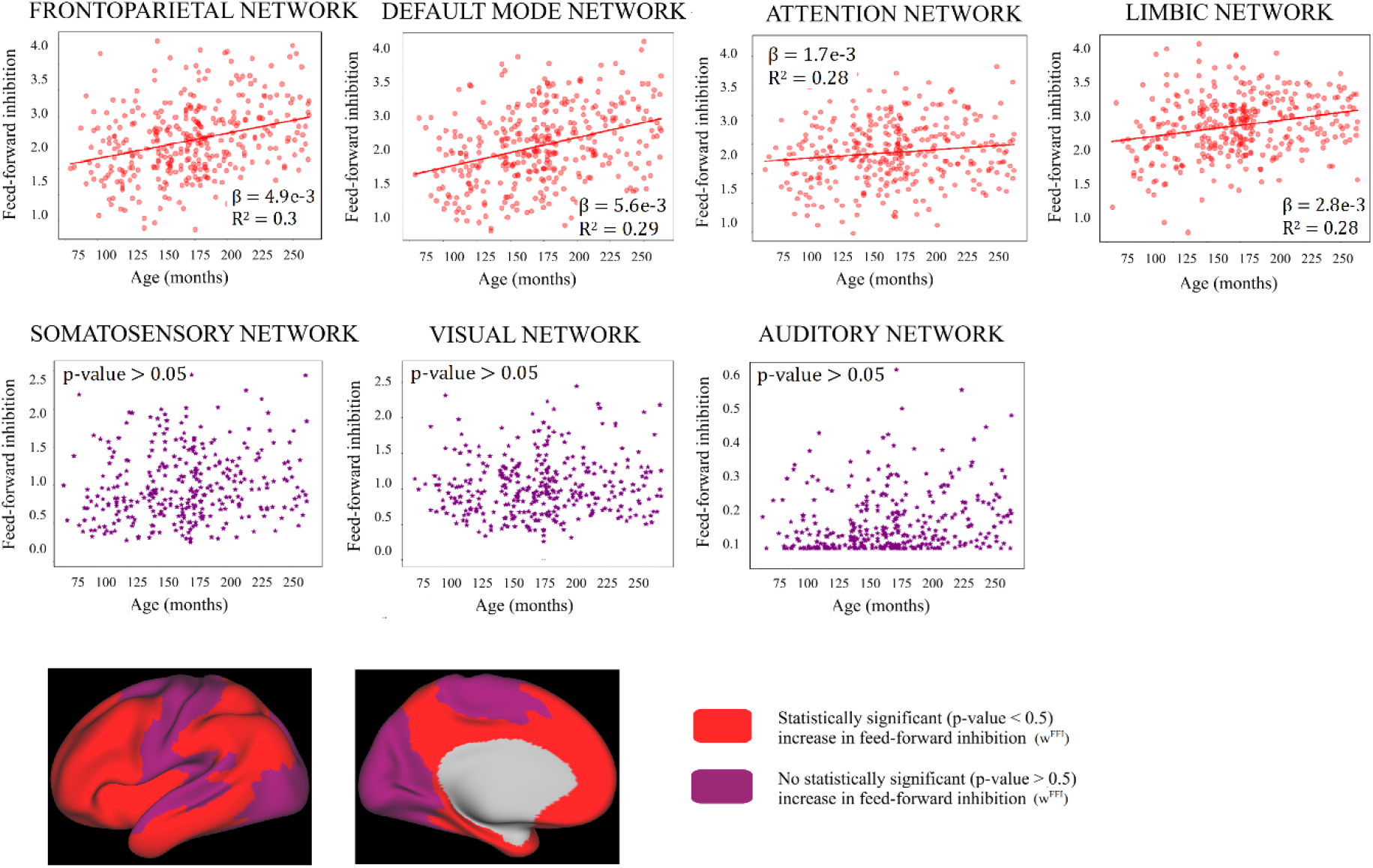
Simulated within-network inhibition levels: Glasser parcels were subdivided into frontoparietal, default mode, attention, limbic, somatosensory, auditory and visual to allow comparison with the empirical group Independent Component/Dual Regression (gIC/DR) findings. Individual data points represent subject-specific parameter 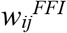 levels within these resting state networks (RSN). Linear regression trajectories (p-value < 0.05) were added alongside correlation coefficient β and determination of coefficient of determination *R*^*2*^. Inflated cortical surfaces illustrate the anatomical distribution of modelled RSNs with and without an age-correlated increase in within-network inhibition levels.

**Figure 10:**
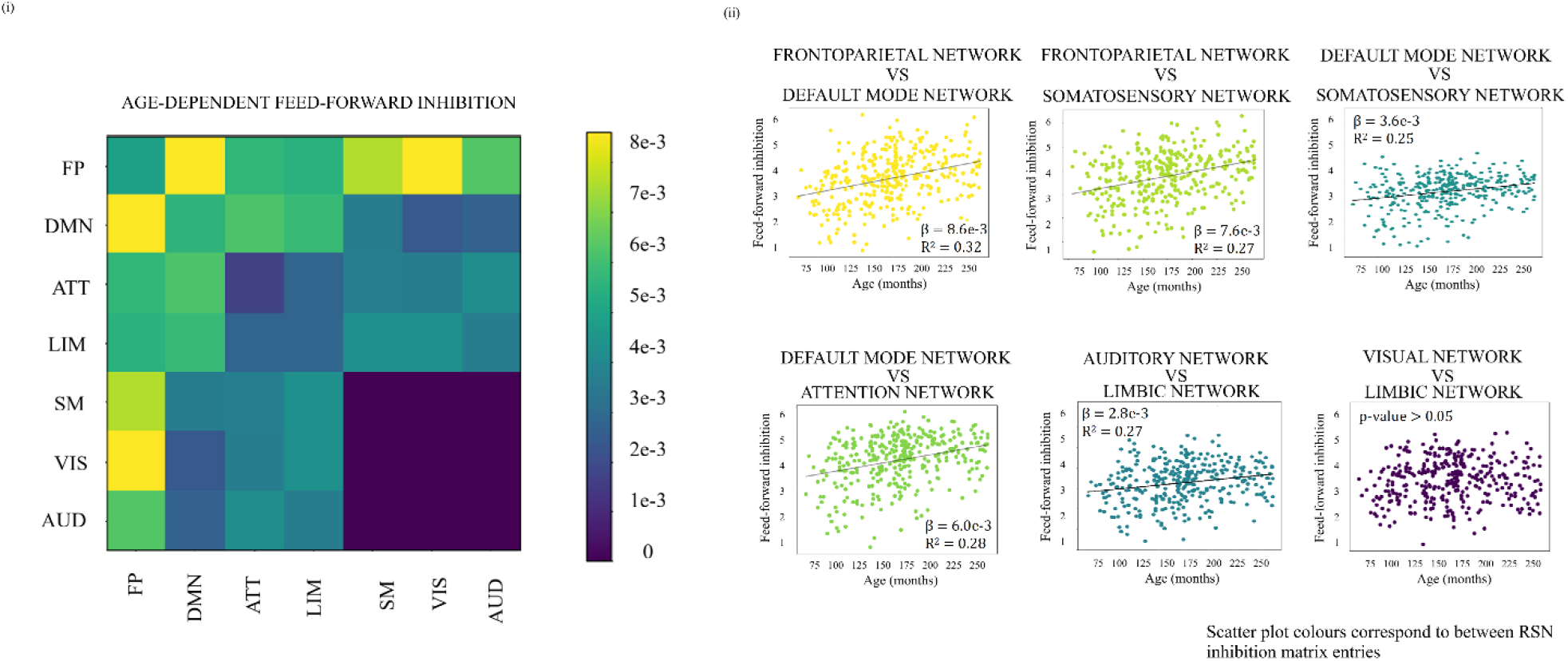
Age dependency of simulated between-RSN feed-forward inhibition levels: **(i):** Matrix entry (*i,j)* represents regression coefficients *β* (p-value < 0.05) of feed-forward inhibition *w*^*FFI*^ across RSNs *(i, j)* regressed against age. **(ii):** representative between-RSN feed-forward inhibition *w*^*FFI*^ regressed against age scatter plots. Regression coefficients *β* correspond to matrix entries in panel (i). *β*, R^2^ presented only for significant regressions.

The model exhibits cross-age RSN-specific trajectories similar to those found in our gICA/DR study. The simulated between-network changes in inhibitory feed-forward coupling followed a similar pattern as empirical between-network FC changes with the most prominent changes inovlving high-order, associational networks, stable primary sensory coupling and attention network connections taking on an intermediate position.

## Discussion

To construct a large-scale computational neuromaturation model capturing age-dependent trajectories in FC development, we first characterized cross-age RSN redistribution throughout childhood and adolescence. We hereafter constructed a computational TVB model allowing region-wise fitting of cortical inhibition. Our empirical study demonstrated cross-age RSN-specific functional and topological maturation. Regional cortical inhibition levels exhibited similar cross-age RSN-dependent trajectories. Thus, our TVB model was not only able to simulate empirical FC but was also able to capture a complex neuromaturational process.

Previous studies have demonstrated that both the adult brain and the brain under development are organized along a functional-structural primary sensory-attention-association axis (Hill et al., 2010; Huntenburg et al., 2018; Sydnor et al., 2021, 2023). Our empirical study consistently demonstrated that large-scale cortical development was differentiated depending on RSN function. High-order, association networks were characterized by topographical redistribution with FP and DMN engaging gradually larger brain areas during childhood and adolescence. The corresponding high-order topology underwent a within-network increase in both degree and strength centrality. Thus, high-order RSNs underwent an age-dependent network integration combined with an increase in information processing, consistent with previous studies of association cortex hub formation (Dong et al., 2020). We also found an association between-network decline in FC strength previously suggested to support efficient processing of cognitive and executive functions (Lopez et al., 2020; Pines et al., 2022; Sherman et al., 2014). Unimodal primary sensory auditory and SM networks remained topographically and topologically stable during the study age range, indicating that these primary sensory regions mature in early childhood. Between-network connectivity changes varied, with both increase and decrease in primary sensory-primary sensory and primary sensory-attention FC strengths. Previous studies have also reported such mixed patterns of combined segregation and integration, potentially supporting goal-driven attention (Keller et al., 2023; Luo et al., 2024; Rohr et al., 2016; Tooley et al., 2022; Wu et al., 2012). It is noteworthy that the visual network topographic distributions followed a different, quadratic trajectory, with the largest cortical representation at early adolescence. Previous studies have similarly demonstrated that visual cortical maturation differs from other primary sensory regions, potentially reflecting a protracted refinement of visual information processing (Luo et al., 2024). Attention networks underwent a reduction in cortical representation and in degree and strength centrality. These findings might reflect an age dependent network pruning and refinement, subserving cognitive and socioemotional development (Dong et al., 2020; Tooley et al., 2022). However, it is previously described that the literature contains inconsistent results on attention network connectivity, with some studies showing an increase in within-network connectivity (Dong et al., 2024, Luo et al., 2024)

The subject specific TVB-Child model reproduced the individual empirical FCs with high fidelity, demonstrating that the model achieved a high level of biorealism. Both within- and between-simulated network age-correlated inhibitory level trajectories varied with the RSN analyzed. FP and DMN networks characterized by age dependent network expansion and network hub emergence as inferred from the empirical data and by model inferred increases of within-network feed-forward inhibition levels. Attention networks underwent age-dependent topographic redistribution reflecting network pruning as of the empirical data paralleled by less pronounced model-inferred increases of feed-forward inhibition. For all primary sensory networks, except for visual network, we did not identify significant network changes through analysis of empirical data or modeling throughout childhood and adolescence. Except, simulated visual networks did not exhibit significant age-related changes of inhibition parameter values, while fMRI data analysis demonstrated topographical redistribution. Comparing empirical between-network FC strength changes with simulated between-network inhibitory couplings revealed similar patterns, with the steepest trajectories seen within high-order regions, low-order regions characterized by no changes and intermediate region trajectories positioned in the middle.

Animal studies and human post-mortem studies have demonstrated that inhibitory upregulation plays an important role in long-term developmental neuromodulation (Bosman et al., 2002; Erzurumlu & Gaspar, 2012; Runge et al., 2020). Exploring such processes *in vivo* in humans is challenging why computational models can provide important insights into the brain dynamics of development. The used brain network model incorporates both FIC, and region-wise tuning of excitation and inhibition following Schirner et al. 2023. We found that the most prominent age-dependent upregulation of feed-forward inhibition occurred within high-order associational networks, that no significant changes occurred in primary sensory networks and that the attention and limbic networks were characterized by intermediate inhibitory upregulation. A small number of human post-mortem studies have demonstrated that human prefrontal cortex undergoes prolonged synaptic reorganization until the third decade of life. On the other hand, primary sensory cortex synaptic density reaches adult levels already during childhood (Huttenlocher & Dabholkar, 1997; Petanjek et al., 2011; Peter R., 1979). Underlying driving neurobiological processes presumably follows same patterns, with maturation of unimodal networks finishing in childhood, and multimodal network maturation continuing throughout adolescence. The computational model similarly revealed that association cross-age RSNs were characterized by the most pronounced inhibitory upregulation, thus reflecting age dependent cortical redistribution. In addition, persistent model inhibition levels are in accordance with childhood completion of primary sensory maturation.

The model provides candidate hypotheses for complex neuromaturational processes demonstrating the potential of dynamic brain network models in improving our understanding of neurodevelopment.

### Limitations

Model simulation outputs need validation in future studies by assessing empirical inhibition levels to confirm validity. Longitudinal MRI GABA spectroscopy studies could provide such data.

## Funding acknowledgement

PR acknowledges supported by the EU Horizon Europe program: BRIDGE (101219311), EBRAINS2.0 (101147319), Virtual Brain Twin (101137289), EBRAINS-PREP 101079717, AISN 101057655, EBRAIN-Health 101058516, EIC grant PHRASE 101058240, the Digital Europe Programme: TEF-Health (101100700), SHAIPED (101195135), CoordinaTEF (101168074), the German Research Foundation: SFB 1436 (project ID 425899996); SFB 1315 (project ID 327654276); SFB 936 (project ID 178316478); SPP Computational Connectomics RI 2073/6-1, RI 2073/10-2, RI 2073/9-1; DFG Clinical Research Group BECAUSE-Y 504745852, DFG German-Spanish project BrainsAge RI 2073/14-1, DFG Project-ID 424778381 - TRR 295 Retune, ERA PerMed JTC2021 German Federal Ministry for Health PatternCog ZMI5-2522FSB904, Berlin University Alliance OpenMake, the Virtual Research Environment at the Charité Berlin and EBRAINS Health Data Cloud and the Berlin Institute of Health and Foundation Charité.

## Notes

### Competing Interest Statement

The authors have declared no competing interest.

